# testCryopreservation of human mucosal leukocytes

**DOI:** 10.1101/039578

**Authors:** Sean M. Hughes, Zhiquan Shu, Claire N. Levy, April L. Ferre, Heather Hartig, Cifeng Fang, Gretchen Lentz, Michael Fialkow, Anna C. Kirby, Kristina M. Adams Waldorf, Ronald S Veazey, Anja Germann, Hagen von Briesen, M. Juliana McElrath, Charlene S. Dezzutti, Elizabeth Sinclair, Chris A. R. Baker, Barbara L Shacklett, Dayong Gao, Florian Hladik

## Abstract

**Background:** Understanding how leukocytes in the cervicovaginal and colorectal mucosae respond to pathogens, and how medical interventions affect these responses, is important for developing better tools to prevent HIV and other sexually transmitted infections. An effective cryopreservation protocol for these cells following their isolation will make studying them more feasible.

**Methods and findings:** To find an optimal cryopreservation protocol for mucosal mononuclear leukocytes, we compared cryopreservation media and procedures using human vaginal leukocytes and confirmed our results with endocervical and colorectal leukocytes. Specifically, we measured the recovery of viable vaginal T cells and macrophages after cryopreservation with different cryopreservation media and handling procedures. We found several cryopreservation media that led to recoveries above 75%. Limiting the number and volume of washes increased the fraction of cells recovered by 10-15%, possibly due to the small cell numbers in mucosal samples. We confirmed that our cryopreservation protocol also works well for both endocervical and colorectal leukocytes.

Cryopreserved leukocytes had slightly increased cytokine responses to antigenic stimulation relative to the same cells tested fresh. Additionally, we tested whether it is better to cryopreserve endocervical cells on the cytobrush or in suspension.

**Conclusions:** Leukocytes from cervicovaginal and colorectal tissues can be cryopreserved with good recovery of functional, viable cells using several different cryopreservation media. The number and volume of washes has an experimentally meaningful effect on the percentage of cells recovered. We provide a detailed, step-by-step protocol with best practices for cryopreservation of mucosal leukocytes.

## Introduction

To develop preventive interventions and therapies for sexually transmitted infections (STIs), it is important to understand how they affect mucosal immunity. Clinical trials of vaccines designed to prevent human immunodeficiency virus (HIV) or herpes simplex virus infection are conducted at sites around the world. Ideally, these trials would include investigation of the cellular immune responses elicited at the mucosal sites where these pathogens initially invade. These analyses require mucosal cell and tissue samples to be stored and shipped to central laboratories, but this is not currently done due to inconsistencies in cryopreservation. Similarly, the effect of topical anti-HIV microbicides on mucosal immunophysiology could be more easily studied if trial sites were able to cryopreserve viable mucosal cell and tissue samples. Thus, little is learned about mucosal cellular immune responses from clinical trials.

While leukocytes isolated from the peripheral blood are routinely cryopreserved for storage and transport, it is currently unclear whether mucosal leukocytes can be cryopreserved successfully [1,2]. Indeed, the fundamental physical characteristics of mucosal leukocytes may differ from those in blood, and as the optimal cryopreservation protocol depends on the physical characteristics of the cells, different protocols may be necessary [3]. In particular, the permeability of cell membranes to water and cryoprotective agents (CPAs) at different temperatures influences the choice of CPA to use in the cryopreservation medium and the rate at which to freeze the cells [3].

We set out to develop an optimal procedure for the cryopreservation of mucosal leukocytes, including formulation of the cryopreservation medium. We isolated T cells and macrophages from the human vagina and measured their physical properties relevant to cryopreservation, as reported previously [4,5]. Based on these measurements, we conducted a series of cryopreservation studies to determine the protocol that leads to maximal recovery of live, functional cells. We subsequently showed that this protocol can be used on cells isolated from endocervical cytobrushes as well as from colorectal biopsies with similar success. A detailed, step-by-step protocol with best practices for cryopreservation of mucosal leukocytes is provided as supporting File S1.

## Methods

### Sample collection

Vaginal tissues discarded as part of vaginal repair surgeries were collected under a waiver of consent at the University of Washington Medical Center (IRB #1167). Endocervical cytobrushes were obtained from volunteers at the University of Washington Medical Center and the Seattle Vaccine Trials Unit (IRB #1830). Rectal biopsies were obtained by flexible sigmoidoscopy from study participants at San Francisco General Hospital (UCSF IRB #627-9 and 569-9). All participants provided written informed consent.

### Cell isolation from vaginal tissue

Vaginal tissue was trimmed free of stroma to a final thickness of 2 mm. The tissue was then minced to pieces smaller than 2x2x2 mm and digested as described previously [6]. Briefly, the tissue pieces were incubated at 37°C for 30 minutes in digestion medium (700 collagen digesting units per mL of collagenase type 2 [catalog number C6885, Sigma-Aldrich] with 500-1000 units per mL DNase I [catalog number DN25, Sigma-Aldrich] in a 1:1 mixture of phosphate buffered saline and R15 [RPMI-1640 containing penicillin-streptomycin and L-glutamine and 15% heat inactivated fetal bovine serum (Gemini Bio-Products, West Sacramento, CA, USA)]). Note that more DNase was used in this procedure than in the reference procedure due to the larger amount of tissue used. After digestion, the tissue was disrupted by ten passages through a 16 gauge needle and isolated cells were recovered through a 70 μm strainer. Digestion was repeated up to three additional times and all cells were combined.

### Cell isolation from endocervical cytobrushes

Cells were isolated from two cytobrushes per donor as described previously [6]. Briefly, the cytobrushes were inserted into the tip of a 25 mL serological pipette filled with phosphate buffered saline. Cells were washed through a 100 μm strainer and collected by gentle expulsion of the saline and movement of the cytobrush in and out of the tip of the pipette. The transport media was also collected and the tube washed out. Cells isolated from the two cytobrushes from the same donor were combined, except where noted below.

### Cell isolation from colorectal biopsies

Rectal biopsies were digested using a procedure similar to that for the vaginal tissue, with the following modifications. Ten biopsies were incubated in digestion medium containing 50 μg/mL Liberase DL (catalog number 05 466 202 001, Roche Diagnostics, Mannheim, Germany), 100U/mL penicillin, 100 μg/mL streptomycin (Gemini Bio-Products), 2 mM L-glutamine (Gemini Bio-Products), 25 mM HEPES buffer (Life Technologies, Grand Island, NY, USA), in RPMI-1640 (Corning/Mediatech, Manassas, VA, USA.)

### Chemicals and cryoprotective agents

Dimethyl sulfoxide (DMSO), ethylene glycol (EG), propylene glycol (PG), glycerol, bovine serum albumin (BSA), Staphylococcal enterotoxin B (SEB), phorbol 12-myristate 13-acetate (PMA), ionomycin, and brefeldin A were all obtained from Sigma-Aldrich. Hydroxyethyl starch 200/0.5 (catalog number V0118) was obtained from AK Scientific (Union City, CA, USA). Trehalose dihydrate and benzonase were obtained from EMD Millipore (Billerica, MA, USA). Cytomegalovirus, Epstein-Barr virus, and influenza peptides (CEF peptide pool) were obtained from AnaSpec (Fremont, CA, USA).

### Cryopreservation procedure

For vaginal cells, 2*10^5^ cells in 200 μL were used per tube (approximately 10% of these were leukocytes) and conditions were tested in duplicate. For cytobrushes, cell suspensions were split among the conditions and tested in singlicate. For colorectal tissues, 1.75-2.5*10^6^ cells in 1 mL per tube were used. There were three procedures for cryopreservation: procedures A and B, which differ in the temperature and volume of the media after thawing, and procedure C, “HANC”, based on the HIV/AIDS Network Coordination blood processing protocol. The latest HANC Cross-Network PBMC Processing SOP is available at https://www.hanc.info/labs/labresources/procedures/Pages/pbmcSop.aspx. **Table 1** summarizes the three cryopreservation procedures.

**Table 1:**
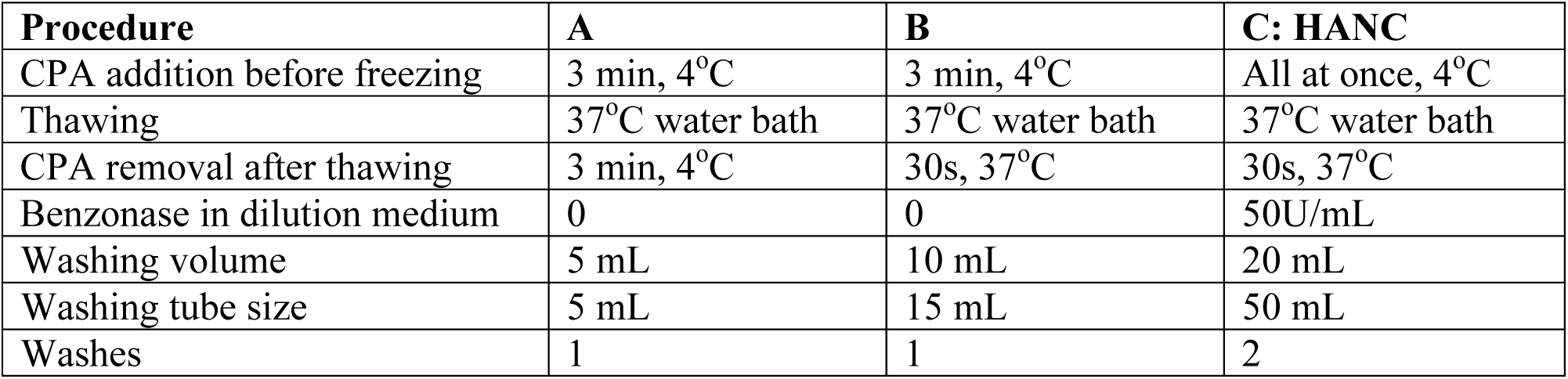
Overview of the three different cryopreservation procedures used.

Cells were suspended in 100 μL of the appropriate cryopreservation medium without any CPAs and aliquoted to cryovials in an ice water bath. 100 μL of a 2X cryopreservation medium (compositions described in Results) was added to each cryovial, dropwise over three minutes for the first two procedures and all at once for the HANC procedure. Cryovials were then transferred to a controlled rate cooling device (Mr. Frosty ‐1°C/min) and frozen in a - 80°C freezer. The cryovials were transferred to the vapor phase of a liquid nitrogen freezer within 3 days, where they were stored for at least one week.

### Thawing

Cells were thawed by quick transfer of the cryovial directly from liquid nitrogen to a 37°C water bath. Once less than a pea-sized amount of ice remained, cryovials were transferred to a biosafety cabinet and opened. Three different procedures were used from this point on, as described in Table 1. For procedure A, cryovials were placed in an ice water bath once the cell suspension was completely thawed and 1.8 mL 4°C cell culture medium was added dropwise over three minutes to dilute the cryoprotective agent(s).

The cells were then transferred to a 12x75 mm polystyrene FACS tube containing three mL precooled cell culture media and washed by centrifugation. For procedure B, no ice bath was used, the cell culture medium was warmed to 37°C, the cell culture medium was added over 30s, and the cells were transferred to a 15 mL polypropylene tube containing 10 mL warmed cell culture medium. The HANC procedure was the same as procedure B, except that 50U/mL benzonase was included in the cell culture medium, the cells were transferred to 20 mL of warmed cell culture medium in a 50 mL polypropylene tube, and the cells were washed twice by centrifugation instead of once. Viability and recovery were measured immediately after thawing, while for intracellular cytokine staining assays, cells were rested overnight prior to stimulation (as described below).

### Flow cytometry

For viability and recovery measurements, cells were stained with fixable live/dead aqua or yellow (Life Technologies, Grand Island, NY, USA) for exactly 20 min, washed, stained with phenotyping markers, washed again, transferred to Trucount absolute counting tubes (BD Biosciences, San Jose, CA, USA), and fixed with 1% paraformaldehyde. Catalog numbers, antibody clones, fluorochromes, and companies are described for all antibodies in supporting File S2. For vaginal cells, phenotyping markers were CD45, CD3, and CD14. For cytobrushes, phenotyping markers were CD45, CD3, CD8, CD14, HLA-DQ, CD19, CD4, and CD66b. All antibodies were titrated before use and used at the minimum saturating dose. Samples were acquired on a five laser BD LSRII (355, 405, 488, 535, and 633 nm) within 1 week.

Absolute counts of rectal cells were obtained using a BD Accuri C6. Cells were stained with CD45, CD4 and CD3 (see supporting File S2) using a no-wash protocol and run directly after staining. Propidium iodide was added just before samples were run to assess viability. Additional phenotyping was performed on rectal cells as described above for genital tract cells, except that Trucount tubes were not used for cell counting. Cells were stained with CD45, CD3, CD4, CD8, CD33, CD66b, CD13, and CD206 (see supporting File S2).

For intracellular cytokine staining, cells were rested overnight in R15 containing 0.5 mg/mL piperacillin/tazobactam (Zosyn, Wyeth-Ayerst, Princeton, NJ, USA). Cells were then stimulated with either pooled peptides from cytomegalovirus, Epstein-Barr virus, and influenza (CEF peptide pool, 3.5 μg/mL each peptide), SEB (5 μg/mL), or PMA (50 ng/mL) with ionomycin (500 ng/mL), with DMSO (peptide vehicle) serving as a negative control. Cells were then incubated for 5 hours at 37°C/5% CO_2_ in the presence of protein transport inhibitors brefeldin A (5 μg/mL) and monensin (GolgiStop, 1 μM), as well as costimulatory antibodies CD28 and CD49d, and degranulation marker CD107a (all from BD Biosciences.)

Following incubation, cells were stained with antibodies for surface markers CD4, CD8, and fixable live/dead aqua viability dye (from Life Technologies). Cells were then fixed in 4% paraformaldehyde and permeabilized using FACS Perm 2 (BD Biosciences.) Intracellular staining was performed with the following fluorochrome-conjugated antibodies: CD3, interferon-γ (IFN-γ), tumor necrosis factor-α (TNF-α), interleukin 2 (IL-2), and macrophage inflammatory protein-1β (MIP-1β) (see supporting File S2 for details about antibodies). Data were acquired within 24 hours on a BD LSRII flow cytometer (405, 488, and 635 nm).

### Data and statistical analysis

Vaginal cells were gated on CD45^+^ and divided into CD14^+^CD3^−^ and CD14^−^CD3^+^. Cytobrushes were additionally gated for CD66b+ (neutrophilic granulocytes). Each population was subsequently gated for viability. Flow cytometric analysis was done in FlowJo version 9 (FlowJo, LLC, Ashland, OR, USA). The number of cells per sample was determined using Trucount tubes which contain a known number of beads. The number of beads acquired was divided by the number of beads known to be in the tube to yield the fraction of the sample acquired. The number of cells of interest was divided by the fraction of sample acquired to yield the number of cells per sample. Samples were acquired at flow rates such that less than five percent of events were aborted.

For rectal cells, absolute counts of CD45 counts were obtained directly using the Accuri. The CD3, CD13, CD33, CD206, and CD66b cells were measured as a percentage of CD45^+^ cells and the absolute counts of these cells determined relative to the absolute counts of the CD45^+^ cells.

Viability is expressed as “absolute viability” or in some cases as “relative viability”, where the viability was normalized to a fresh control sample from the same donor. Recovery reflects the number of live cells of interest recovered as a percentage of the number of live cells of interest in the fresh control sample from the same donor.

To analyze cytokine response data, we employed a previously described statistical test to determine whether antigen-specific or polyclonal responses differed significantly from background responses [7]. This formula assumes a Poisson distribution and takes into account the actual number of gated events as opposed to percentages. Net responses were calculated by subtracting negative control values from stimulated values.

All statistical tests used are indicated in the supplementary tables (supporting File S3. Single comparisons were generally done with paired t-tests and multiple comparisons with repeated measures ANOVA and Tukey post-tests. Data analysis was done using R version 3.2.3 [8] and the packages dplyr [9], magrittr [10], reshape2 [11], tidyr [12], readr [13], RColorBrewer [14,15], nlme [16], multcomp [17], and Hmisc [18]. Plots were created using ggplot2 [19] and tables were rendered with pander [20] and knitr [21,22]. All code and source data are included (supporting File S4).

## Results

### Development of novel cryopreservation medium

We set out to develop an optimal cryopreservation medium for mucosal mononuclear leukocytes. First, we did a series of exploratory experiments, comparing different cryoprotective chemicals and concentrations, with the goal of finding one or two cryopreservation media for more rigorous testing. Because of the exploratory nature of these experiments and the limited amount of replication, no statistical tests were performed in this section. For all of these experiments, cells were cryopreserved using procedure A (Table 1).

Based on measurements of the cryobiological properties of vaginal T cells and macrophages [4,5], we hypothesized that dimethyl sulfoxide (DMSO), ethylene glycol (EG), and propylene glycol (PG) would be effective cryoprotective agents (CPA) for these cells, whereas glycerol would not be due to very limited cell membrane permeability to glycerol. Indeed, cryopreservation with 1.5M glycerol in FBS leads to considerably worse viability than either 10% DMSO or 1.5M PG for both T cells and macrophages (**Figure 1A)**.

**Figure 1.**
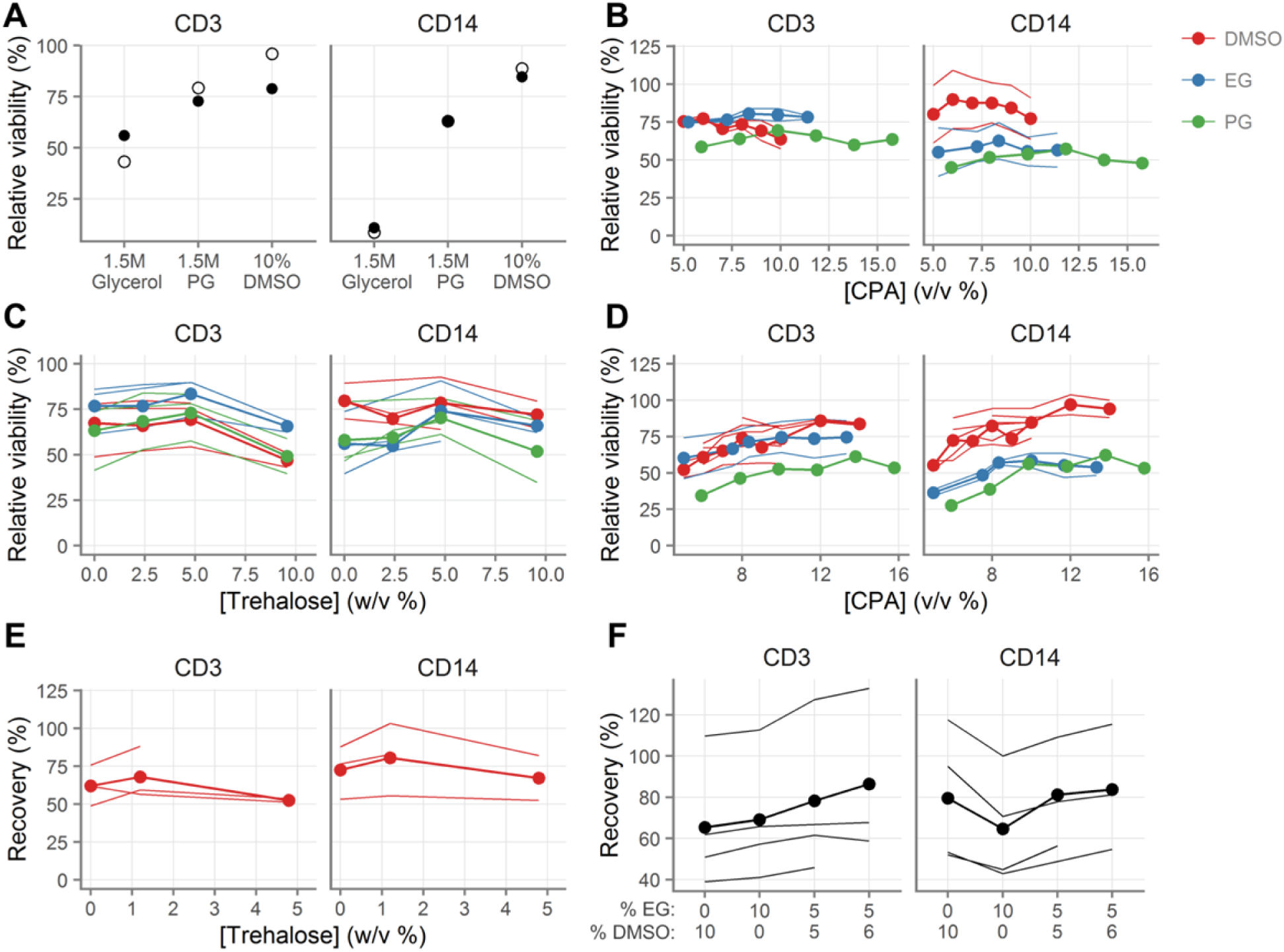
Development of novel cryopreservation medium. **A**, Viability of vaginal T cells (indicated “CD3”) and macrophages (indicated “CD14”) following cryopreservation with 1.5M glycerol, 1.5M propylene glycol (PG) or 10% dimethyl sulfoxide (DMSO). **B**, Effect of CPA concentration on cell viability after cryopreservation. **C**, Effect of trehalose concentration on cell viability after cryopreservation in the presence 0.84M DMSO, 1.5M PG, or 1.75M EG. **D**, Effect of CPA concentration on cell viability after cryopreservation in the presence of 4.8% trehalose. **E**, Effect of trehalose concentration on recovery after cryopreservation in the presence of 1.4M DMSO. **F**, Effect of mixtures of DMSO and EG on cell recovery after cryopreservation in the presence of 1.2% trehalose. Each line for a given color indicates a different tissue sample. The line with the circles indicates the mean across the different tissue samples. Measurements were made in duplicate for each tissue sample. For relative viability, 100% indicates the same viability as the fresh cells.

We had previously determined CPA toxicity on mucosal cells without freezing and thawing, by adding DMSO, EG, or PG at various concentrations dropwise to cell suspensions, incubating for ten minutes on ice, and then dropwise diluting them out. At concentrations below 10% v/v, all three chemicals had a minor negative impact on cell viability, of less than 10% at the highest concentrations [4]. We then tested their cryoprotective effectiveness by measuring the viability of cells after cryopreservation (**Figure 1B**). T cells were preserved similarly with all three CPAs, whereas macrophages appeared to be preserved better with DMSO.

DMSO, PG, and EG are permeating CPAs, meaning that they displace water from the cell and prevent cell lysis. Combining permeating CPAs with nonpermeating CPAs (commonly sugars and polymers), which have a different mechanism of cryoprotection, can improve recovery [23–25]. We therefore cryopreserved cells in the presence of permeating CPAs and a range of concentrations of the nonpermeating CPA trehalose, a non-cell permeant disaccharide, and found moderately better viability than without trehalose (**Figure 1C**). We then re-titrated the intracellular CPAs in the presence of trehalose. These experiments suggested that 10-12% (v/v) was optimal for all intracellular CPAs in the presence of 4.8% (w/v) trehalose (**Figure 1D**).

Finally, we tested trehalose again in the presence of DMSO, this time measuring recovery of live cells after cryopreservation rather than just viability, and found that 1.2% trehalose appeared equivalent to 4.8% (**Figure 1E**).

We hypothesized that a combination of EG and DMSO in the presence of 1.2% trehalose might be better than either alone. We tested several combinations, and found that 5% EG with 6% DMSO seemed superior for absolute recovery (**Figure 1F**) than either EG or DMSO alone.

### Validation of novel cryopreservation medium

Based on these exploratory experiments, we selected three cryopreservation media to formally compare: 10% DMSO, 8% DMSO with 1.2% trehalose, and 6% DMSO + 5% EG with 1.2% trehalose (all v/v, except trehalose w/v, all diluted in FBS). Samples from ten donors were cryopreserved with procedure A (Table 1). in duplicate for each condition and both viability and recovery were measured.

2. The results of this experiment are summarized in **Figure 2A–B**; all comparisons and effect sizes are detailed in **Tables S1-8**. Overall, the three-CPA cocktail was best for T cells and no different from 10% DMSO for macrophages. The DMSO and trehalose mixture was better than 10% DMSO for both cell types, but not as good as the three-CPA cocktail for T cells. The highest recovery for T cells was with the three-CPA cocktail (68.0%, [95% CI: 56.4, 79.6]). The highest recovery for macrophages was with the DMSO and trehalose mixture (72.9% [63.0, 82.8]). Recovery with the three-CPA cocktail was fifteen percentage points greater than with 10% DMSO (p < 0.0001) for T cells. Recovery with the DMSO and trehalose mixture was about five percentage points greater than with 10% DMSO (p = 0.06) for macrophages. Of note, for T cells but not for macrophages, recovery was predicted by cell viability (**Figure 2C, Table S9**).

**Figure 2.**
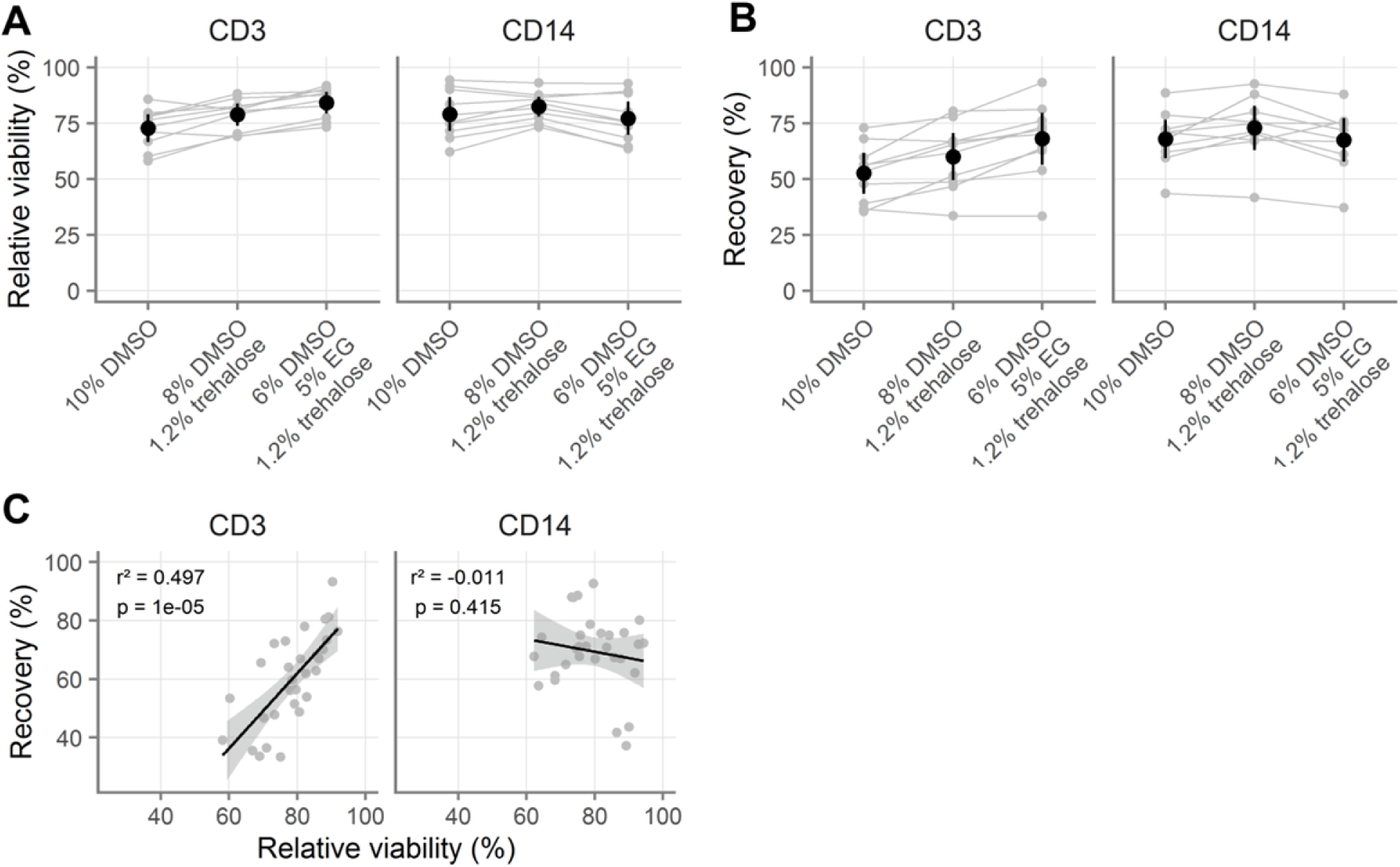
Validation of novel cryopreservation medium. A-B, Comparison of three cryopreservation media by relative viability (A) and recovery (B) for vaginal T cells (indicated “CD3”) and macrophages (indicated “CD14”). All samples were processed with procedure A (see Table 1). **C**, Correlation of relative viability and recovery for vaginal T cells and macrophages with different cryomedia. For relative viability, 100% indicates the same viability as the fresh cells. Gray symbols indicate the average of duplicates from the same sample, with gray lines indicating pairing. Black symbols show the mean across all samples and black vertical lines show the 95% confidence interval of the mean. In C, black line indicates the slope of the linear model and the grey shaded area indicates the 95% confidence interval. Adjusted p-values and r^2^ values are shown from linear regression.

### Comparison with published cryopreservation methods

Having found that 6% DMSO and 5% EG with 1.2% trehalose was superior to 10% DMSO for T cells, we compared it to other cryopreservation media reported in the literature. For these experiments, we switched from using cryopreservation procedure A to procedure B (Table 1) for consistency with published methods. As shown in **Figure S1A** and **Table S10**, there was no difference in cell recovery between procedures A and B. Additionally, we replaced trehalose with 6% hydroxyethyl starch (HES), given a recent report of its benefits for human pluripotent stem cells [26]. Tested with vaginal cells from four donors, we found HES to be superior to trehalose (**Figure S1B, Table S11**), with an 8.66% [95% CI: 3.24, 14.07] increase in recovery for T cells (p = 0.009) and a 7.76% [3.61, 11.91] increase for macrophages (p = 0.01). Consequently, we replaced trehalose with 6% HES for the rest of our experiments.

At this point, we compared our medium to several other published cryopreservation media. We compared it to two serum-free cryomedia developed for PBMC cryopreservation: 10% DMSO alone (GHRC I [Global HIV Vaccine Research Cryorepository]) or 5% DMSO with 6% HES (GHRC II), both in RPMI with 12.5% BSA [27]. We also compared it to the standard cryopreservation media of 10% DMSO in FBS, frozen according to procedure C published by the HIV/AIDS Network Coordination (HANC) (Table 1). All conditions were measured in duplicate with 200,000 cells per cryovial from six tissue donors, as described above. Our cryopreservation medium and the two serum-free media were processed with procedure B.

As shown in **Figure 3A** and **Tables S12-17**, our medium and both GHRC media were better than 10% DMSO processed according to the HANC procedure (difference in mean percent recovered cells: 10.06% to 12.98% for T cells; 13.57% to 17.47% for macrophages; p-values < 0.0001 for all three media for both cell types). However, there was no difference between the GHRC media and our medium (difference in mean percentage of recovered cells: - 0.09% to 2.83% for T cells, p-values > 0.64; ‐1.22% to 2.67% for macrophages, p-values > 0.70). Thus, our cryopreservation medium resulted in similar levels of cell recovery to two serumfree cryopreservatives, and all three were superior to 10% DMSO when the latter was processed according to the procedure C (HANC).

**Figure 3.**
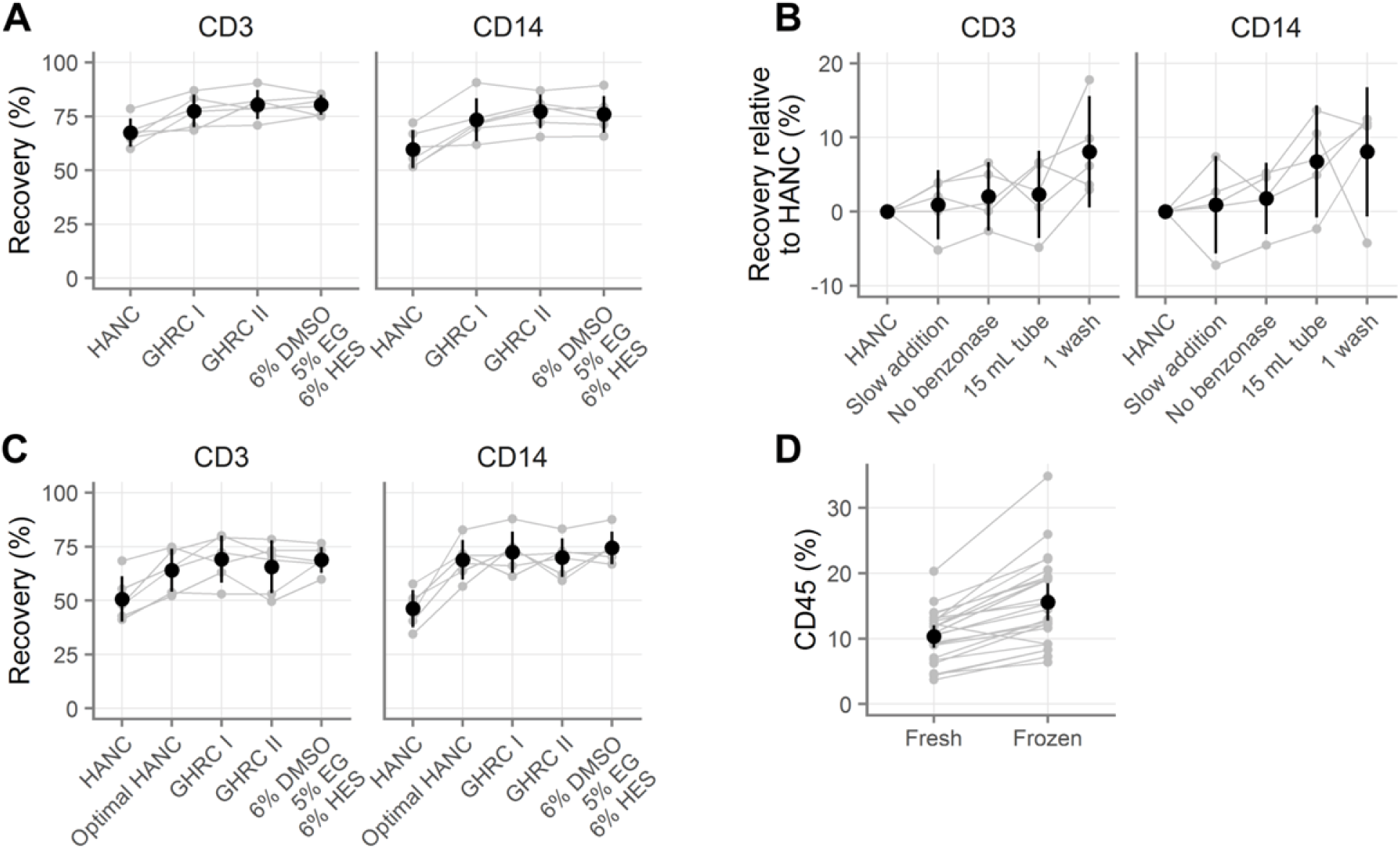
Comparison of novel cryopreservation medium with published methods. **A**, Recovery of vaginal T cells (indicated “CD3”) and macrophages (indicated “CD14”) with several cryopreservation media. GHRC I [Global HIV Vaccine Research Cryorepository] is 10% DMSO alone and GHRC II is 5% DMSO with 6% hydroxyethyl starch, both in RPMI with 12.5% bovine serum albumin. HANC is 10% DMSO in FBS. HANC was processed according to procedure C and the others were processed with procedure B (see Table 1). **B**, Evaluation of which processing steps improve the recovery obtained relative to the standard HANC procedure, where positive values indicate improvements over HANC. **C**, Rescue of HANC procedure (“optimal HANC”) by use of optimal processing procedures for small sample sizes. **D**, Percentage of all vaginal cells that are CD45^+^ before and after freezing. Gray symbols indicate the average of duplicates, with gray lines indicating pairing. Black symbols show the mean across all samples and black vertical lines show the 95% confidence interval of the mean.

Given the success of the GHRC serum-free media, we tested our cryopreservation medium suspended in FBS as usual or in RPMI containing 12.5% bovine serum albumin as in the GHRC media. In nine samples, the choice of diluent did not affect T cell recovery (**Figure S1C** and **Table S18**, 2.04% [95% CI: ‐7.21, 3.13] percent difference in recovery, p = 0.39), but FBS was better for macrophages (5.93% [0.98, 10.89], p = 0.02).

### Optimal processing

It was unclear from the experiment described in Figure 3A whether the difference in recovery was due to differences in processing or differences in the cryopreservation media. Indeed, it seemed somewhat improbable that such a large difference would exist between the 10% DMSO in FBS condition (HANC) and the 10% DMSO in RPMI with 12.5% BSA condition (GHRC I). To determine whether differences in cryopreservation processing or media composition underlie the differences in cell recovery, we systematically tested the variables.

We cryopreserved cells from five donors using the standard HANC protocol, as well as with each of the following modifications: dropwise CPA addition over three minutes, no benzonase, thawing into a 15 mL tube with 10 mL media instead of a 50 mL tube with 20 mL media, or one wash instead of two. **Figure 3B** summarizes the results of that experiment and **Tables S19-26** detail all findings. We found that doing only one wash caused an 8% increase in recovery for both cell types (p = 0.002 and p = 0.05), while using a 15 mL tube increased the recovery by 2.30% for T cells and 6.76% for macrophages (p = 0.82 and p = 0.16). Slow addition of CPA and omitting benzonase did not affect the recovery.

Building on this result, we repeated the experiment in Figure 3A with six more donors, this time including an additional condition of HANC processed optimally (with all four modifications described above; i.e. the same procedure as used for the other cryopreservation media). The results of this experiment show that optimal processing rescues the 10% DMSO in FBS cryopreservation medium (**Figure 3C** and **Tables S27-34**). As before, the HANC condition led to notably lower viability and recovery of both cell types. However, when processed optimally, the recovery was not different than that of the other CPAs.

Additionally, we utilized the large number of samples (n = 24) from these experiments to compare the percentages of scatter-gated events that were live CD45^+^ leukocytes before and after freezing (**Figure 3D, Tables S35-36**). On average, 10.32% [95% CI: 8.61, 12.03] were CD45^+^ before freezing, while 15.59% [12.75, 18.42] were CD45^+^ after freezing (p < 0.0001), indicating a preferential reduction of non-leukocytes during cryopreservation Taken together, these results suggest that for small numbers of mucosal cells (200,000 cells per cryovial), experimentally significant numbers of cells can be saved by limiting washes and using smaller tubes. Additionally, they suggest that any one of several cryopreservation media can be used to obtain recoveries of about 75% for both T cells and macrophages.

### Cryopreservation of endocervical cytobrushes

The previous studies were done with vaginal cells with duplicate cryovials of 200,000 cells each per condition. To determine whether our cryopreservation medium worked well with other mucosal samples and cell types, we obtained two sequential endocervical cytobrushes from thirteen volunteers. Cells from each pair of cytobrushes were isolated and combined. They were then divided evenly by volume and stained fresh or cryopreserved with 6% DMSO, 5% EG and 1.2% trehalose, or with 10% DMSO, both in FBS, using procedure A (Table 1). Due to the limited and variable cell yield inherent to this type of sample, each condition was done in singlicate and the total number of cells per condition varied from donor to donor.

There were no major differences between the two cryopreservation media for the cryopreservation of endocervical cells (**Figure 4A** and **Tables S37-38**). We recovered a mean of 60.87% T cells [95% CI: 37.81, 83.94] with our cryopreservation medium and 53.31% [37.65, 68.96] with 10% DMSO (p = 0.25). For macrophages, we recovered 66.22% [41.65, 90.80] with our cryopreservation medium and 67.74% [39.20, 96.28] with 10% DMSO (p = 0.73). Because cytobrushes also contain large numbers of neutrophils, we measured the recovery of CD66b+ neutrophils. Only 31.20% of live neutrophils [19.70, 42.70] were recovered with our medium and 36.15% [11.48, 60.81] with 10% DMSO (p = 0.67). To determine the average yield of a pair of cytobrushes after cryopreservation, we multiplied the number of cells we recovered after cryopreservation by the number of ways the sample had been divided (i.e. by three if the sample had been stained fresh and cryopreserved with both cryopreservation media).

**Figure 4.**
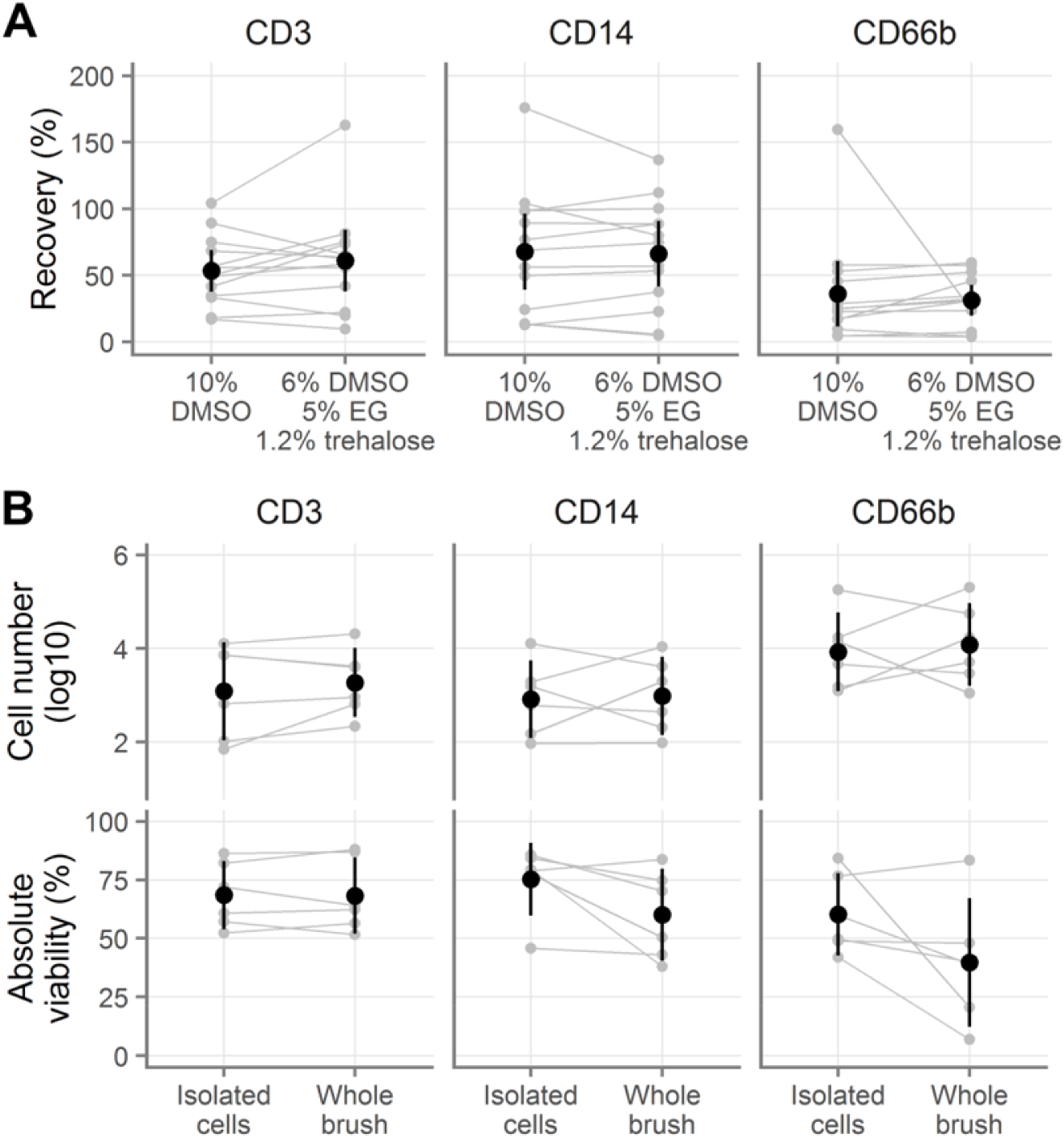
Cryopreservation of cells derived from endocervical cytobrushes. **A**, Recovery of endocervical T cells (indicated “CD3”), macrophages (indicated “CD14”), and neutrophils (indicated “CD66b”), when cryopreserved after isolation from cytobrushes. **B**, Absolute number of live (top) and viability of total endocervical cells (bottom) after cryopreservation either as a cell suspension or still on the cytobrush. Gray symbols indicate the individual samples, with gray lines indicating pairing. Black symbols show the mean across all samples and black vertical lines show the 95% confidence interval of the mean.

The geometric mean numbers of cells yielded from a pair of cytobrushes after cryopreservation was almost identical across the two conditions, with 8,000 macrophages, 4,000 T cells, and 181,000 neutrophils (**Table S39**).

Because isolating the cells from cytobrushes can be inconvenient at clinical trial sites, it would be ideal if cytobrushes could be frozen intact, without removing the cells. To test this possibility, we obtained pairs of cytobrushes from six female volunteers and cryopreserved with 6% DMSO, 5% EG, and 6% HES. We isolated the cells from one cytobrush and cryopreserved the cell suspension. We put the other cytobrush directly into a 4 mL cryovial containing 0.75 mL RPMI with 12.5% BSA. We then added 0.75 mL of 2X cryopreservation medium (i.e. 10% EG, 12% DMSO, and 12% HES), mixed, and cryopreserved the cytobrush. After thawing with procedure B (see Table 1), we isolated the cells from the cytobrush and compared the viability and total live cell numbers. In half of the cases, the first cytobrush taken was cryopreserved intact and in the other half the second was cryopreserved intact.

Similar numbers of live cells of all three cell types were recovered whether frozen in suspension or on the brush (**Figure 4B, top; Table S40-41**), with small effect sizes and all p-values > 0.35. In contrast, there was a difference in viability depending on whether cells were isolated before or after freezing: T cells retained the same viability whether frozen in suspension or on the brush, but macrophages and neutrophils did not (**Figure 4B, bottom; Table S42-43**). The average difference in viability for T cells was less than 1% (p = 0.93). Freezing on the brush rather than in suspension decreased the viability of both CD14 macrophages (by 15.22 percentage points [-2.53, 32.97], p = 0.08) and CD66b neutrophils (by 20.59 percentage points [-6.40, 47.58], p = 0.11).

### Colorectal cells

We also wanted to explore whether cells isolated from other mucosal surfaces can be cryopreserved. To address this question, we isolated cells from the colorectal mucosa from four donors and repeated the experiment described in Figure 3C, freezing 1.75 to 2.5*10^6^ viable leukocytes per cryovial. As shown in **Figure 5A** and **Tables S44-5**, similar numbers of CD3 T cells were recovered from all conditions except for the 6% DMSO, 5% EG, and 6% HES condition, where the recovery was about 10% less (all p < 0.0001). The same was true of CD13 macrophages (**Tables S44, 46**, with three p-values < 0.05 and one p = 0.14). Recovery of CD66b neutrophils was, as expected, poor with all methods (**Figure 5A**). CD33 and CD206 myeloid cells showed similar patterns, though these cells were relatively infrequent (**Table S44**).

**Figure 5.**
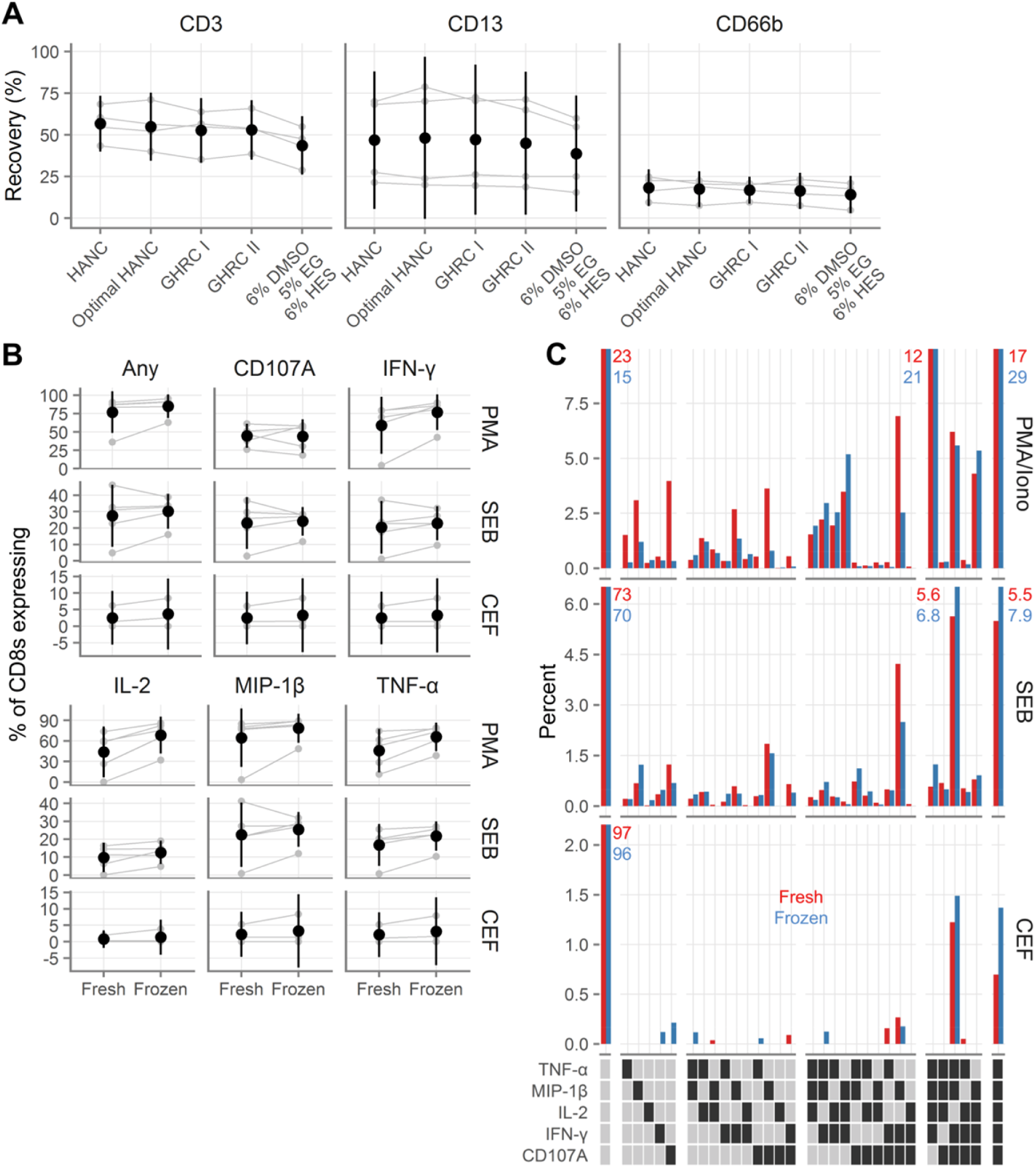
Recovery and functionality of colorectal cells after cryopreservation. **A**, Recovery of colorectal T cells (indicated “CD3”), macrophages (indicated “CD13”), and neutrophils (indicated “CD66b”), after cryopreservation with several cryopreservation media. GHRC I [Global HIV Vaccine Research Cryorepository] is 10% DMSO alone and GHRC II is 5% DMSO with 6% hydroxyethyl starch, both in RPMI with 12.5% bovine serum albumin. HANC is 10% DMSO in FBS. HANC was processed according to procedure C and the others were processed with procedure B (see Table 1). **B**, Background-subtracted cytokine production from colorectal CD8 T cells fresh or after cryopreservation after stimulation with phorbol 12-myristate 13-acetate (PMA); staphylococcal enterotoxin B (SEB); or human cytomegalovirus, Epstein-Barr virus and influenza virus peptides (CEF). Gray symbols indicate the individual samples, with gray lines indicating pairing. Black symbols show the mean across all samples and black vertical lines show the 95% confidence interval of the mean. **C**, Polyfunctionality of colorectal CD8 cytokine responses to stimulation. Red bars indicate fresh samples and blue bars indicate samples tested after cryopreservation. Each bar corresponds to the percent of CD8 cells that expressed the indicated cytokines, averaged across the donors. Percentages are indicated explicitly for bars that exceed the axis limits of the graphs. In the legend, a dark square indicates presence of that cytokine and a light square its absence. White gaps divide the graphs and legends into groups expressing 0,1,2, 3, 4, or 5 cytokines.

Additionally, we tested the functional capacity of rectal cells from 6 donors cryopreserved with 10% DMSO in FBS. After thawing, the cells were rested overnight in media in a 37%C incubator and then we measured cytokine production in response to Staphylococcal enterotoxin B (SEB), phorbol myristate acetate (PMA) with ionomycin, or a pool of peptides from cytomegalovirus, Epstein-Barr virus, and influenza. Percent cytokine-producing CD8^+^ T cells were generally higher in the cryopreserved conditions than in the fresh conditions, though the confidence intervals often overlapped with zero (**Figure 5B, Table S47**). This indicates that cryopreserved rectal immune cells respond as or more robustly to both antigen-specific and superantigen and phorbol ester stimulation as do their fresh counterparts.

We furthermore assessed the polyfunctionality of the responses, comparing fresh to frozen (**Figure 5C**). There was generally little difference between the quality of the responses. In fresh samples, cytokine-producing cells tended to have less polyfunctional responses, while in frozen samples, cytokine-producing cells tended to be slightly more polyfunctional, consistent with slightly increased cytokine responses in cryopreserved samples.

## Discussion

To develop preventive interventions for HIV and other STIs, we need to understand how leukocytes in the cervicovaginal and colorectal mucosae respond to pathogens and the interventions themselves. Gaining this knowledge is complicated by the lack of an effective cryopreservation protocol for mucosal cells. This is especially the case with regard to HIV, because of the need to ship samples to central laboratories from trial sites in sub-Saharan Africa where HIV is highly prevalent and levels of mucosal immune activation tend to be elevated while vaccine responses tend to be muted [28,29].

In this paper, we show that it is possible to cryopreserve small numbers of isolated cervicovaginal and colorectal immune cells and achieve viable cell recoveries of greater than 75%. With colorectal cells we also demonstrate good retention of cellular functionality, measured by stimulated cytokine secretion, after cryopreservation. Due to limited cell numbers, vaginal and endocervical cells were not tested for functionality. We further show that the procedure used to cryopreserve and thaw the cells meaningfully influences recovery, with the number and volume of washes playing the biggest role. Based on our findings, we provide a detailed procedure for cryopreserving isolated mucosal cells in supporting File S1. In a separate, forthcoming manuscript, we will report on cryopreserving whole mucosal tissue specimens. Together, these works will provide guidance on optimal procedures and expected yields.

Our data indicate that several cryopreservation media are largely equivalent: 10% DMSO in FBS or in RPMI with 12.5% BSA, 5% DMSO with 6% HES in RPMI with 12.5% BSA, or 6% DMSO with 5% EG and 6% HES in FBS. It is worth noting that the recovery of colorectal T cells and macrophages was about ten percent lower with the last of these cryomedia than with the others, suggesting that it is suboptimal for immune cells isolated from that tissue. Overall, the equivalence of these media affords investigators flexibility in choosing the medium that has the chemical characteristics most suited to their needs. For example, for some applications serum-free cryopreservation media or a lower than 10% DMSO concentration may be desirable.

The serum-free cryopreservation media used here have been used with PBMC in several publications. One group reported lower recovery and higher background in ELISPOT assays using these media than with serum-containing or other serum-free media [30]. However, two other publications report similar or superior recovery and background with these media in comparison to cryopreservation media containing FBS [27,31]. Investigators should therefore test their specific application.

We found that an additional wash in a larger volume caused a substantial decrease in recovery of vaginal immune cells. However, this was not the case with colorectal cells, where the recoveries were equivalent. The discrepancy between the vaginal and colorectal recoveries may be a result of differences in cell numbers. For the vaginal samples, 200,000 total cells, including 10-20,000 leukocytes, were used in each cryovial as this is the expected yield of endocervical cytobrushes. Ten times as many leukocytes were frozen per cryovial in the colorectal cases. Loss during washing may be less noticeable with larger cell numbers, as indeed is suggested by the regular use of the two-wash procedure by member groups of the Office of HIV-AIDS Network Coordination (HANC) for PBMCs frozen at 10-20*10^6^ per vial with excellent recovery [32].

Somewhat surprisingly, a greater fraction of cryopreserved colorectal cells produced cytokines in response to antigen or superantigen than did their fresh counterparts (Figure 5B). In addition, we saw a consistent enrichment of CD45-expressing cells after freezing (Figure 3D). This may indicate that cryopreservation with the media used here preferentially preserves mucosal leukocytes over other cell types. The increase in leukocytes as a fraction of all cells could cause increased cytokine production by improving the ability of antigen-presenting cells to find T cells or decreasing the potential inhibitory activity of other contaminating cell types. Alternatively, the process of cryopreservation itself or a higher presence of dead cells in thawed compared to fresh samples could also cause non-specific immune activation.

Additionally, our data suggest that it may be possible to cryopreserve endocervical cells without removing them from the cytobrush first. Similar numbers of live T cells, macrophages, and neutrophils were recovered whether the cells were isolated from the brush before or after cryopreservation. However, the viability of macrophages and neutrophils, but not T cells, was reduced when cryopreserved on the brush, so more testing is needed.

In conclusion, we have shown several cryopreservation media that can be used to obtain good recoveries of genital and colorectal leukocytes. Additionally, we have shown that these cells have similar or greater functionality than similar cells processed fresh. Based on our results, we provide a detailed protocol for optimal cryopreservation of mucosal leukocytes in supporting File S1.

All samples used in this study were stored in liquid nitrogen vapor for short periods of time, between one week and one month, but in practice samples may be stored for years to decades. While we did not formally demonstrate that equivalent cell recovery would result regardless of the length of storage, several other groups have. In the context of blood donation, equal recovery of white blood cells was shown for samples stored from as briefly as one day to longer than six years [33]. Microarray analysis of these samples did not indicate an effect of storage length. Similarly, hematopoietic progenitor cells showed equivalent viability, time to platelet engraftment, and time to white blood cell engraftment whether stored for less than one year, one to nine years, or more than nine years [34]. Given these findings, we expect that the recoveries attained in our study will generalize to samples stored for longer duration.

In conclusion, we have shown several cryopreservation media that can be used to obtain good recoveries of genital and colorectal leukocytes. Additionally, we have shown that these cells have similar or greater functionality than similar cells processed fresh. Based on our results, we provide a detailed protocol for optimal cryopreservation of mucosal leukocytes in supporting File S1.

## Acknowledgements

We thank the participants for generously making this research possible. We acknowledge everyone at the Seattle VTU clinic and Amy Gest at UW OB/GYN for recruiting participants and obtaining high quality samples. We thank Katharine Westerberg for assistance with processing vaginal tissues. We also thank Dr. Peter Hunt and the San Francisco General Hospital (SFGH) SCOPE investigators and study coordinators for recruiting participants for gastrointestinal biopsies, and Dr. Ma Somsouk and his team at SFGH for obtaining rectal biopsy tissue. We are grateful to Dr. Patricia D’Souza (DAIDS, NIH) for support and encouragement. We thank the James B. Pendleton Charitable Trust for their generous equipment donation. This publication was made possible with help from the University of California San Francisco-Gladstone Institute of Virology & Immunology Center for AIDS Research (CFAR), a National Institutes of Health-funded program (http://www.nih.gov/; P30 AI027763). Funding was provided by the Bill & Melinda Gates Foundation (http://www.gatesfoundation.org/; OPP1032522 to DG and FH) and the National Institutes of Health (Supplement to R33 AI0094412 to FH and DG, HVTN Mucosal Immunology Group Award to FH (parent award: UM1 AI068618 to MJM), and Administrative Supplement to R01 AI057020 to BLS).

## Supporting Information Captions

**Figure S1.**
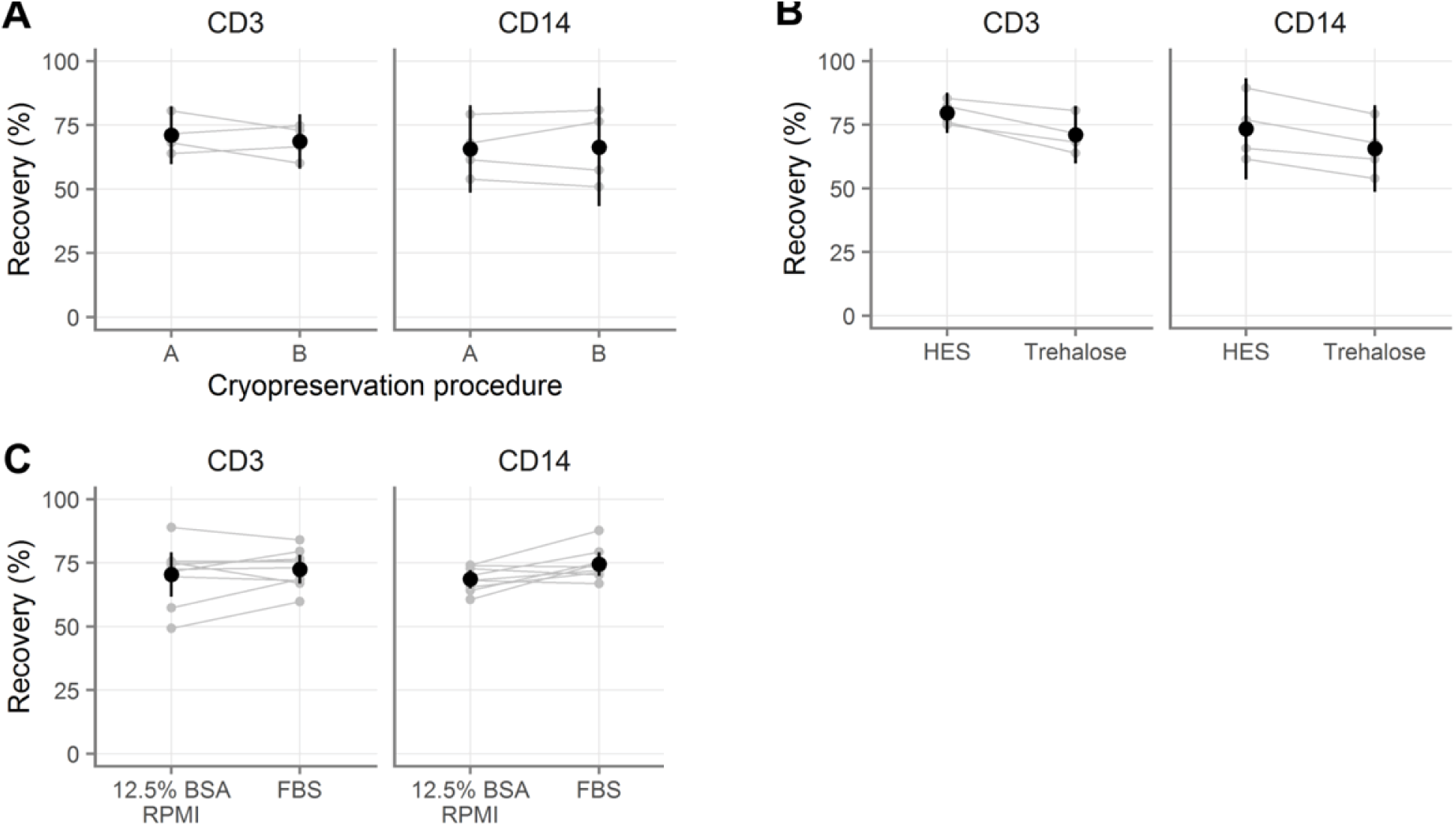
Improvements to cryomedia and processing procedures. **A**, Comparison of different procedures (see Table 1) for cryopreserving vaginal T cells (indicated “CD3”) and macrophages (indicated “CD14”). **B**, Comparison of supplementing 5% EG and 6% DMSO in FBS with 6% HES or 1.2% trehalose. **C**, Comparison of cryopreservation with 5% EG, 6% DMSO, and 6% HES in FBS or in 12.5% BSA in RPMI. Gray symbols indicate the average of duplicates, with gray lines indicating pairing. Black symbols show the mean across all samples and black vertical lines show the 95% confidence interval of the mean.

**S1. Text. Detailed, step-by-step protocol for cryopreservation of mucosal leukocytes.**

**S2. Table. Sourcing information for antibodies used.**

**S3. Statistics. Complete statistical tables.**

**S4. Code and data. Analysis code and raw data files.** Contains the complete R code and raw data needed to reproduce the analyses reported here, as well as generate all figures.

## References

1. Liebenberg LJ, Gamieldien H, Mkhize NN, Jaumdally SZ, Gumbi PP, Denny L, et al. Stability and transport of cervical cytobrushes for isolation of mononuclear cells from the female genital tract. J Immunol Methods. Elsevier B.V.; 2011;367: 47–55. doi:10.1016/j.jim.2011.01.013

2. Bull M, Lee D, Stucky J, Chiu Y-L, Rubin A, Horton H, et al. Defining blood processing parameters for optimal detection of cryopreserved antigen-specific responses for HIV vaccine trials. J Immunol Methods. 2007;322: 57–69. doi:10.1016/j.jim.2007.02.003

3. Mazur P. Kinetics of Water Loss From Cells At Subzero Temperatures and the Likelihood of Intracellular Freezing. J Gen Physiol. 1963;47: 347–369. doi:10.1085/jgp.47.2.347

4. Shu Z, Hughes SM, Fang C, Huang J, Fu B, Zhao G, et al. A Study of the Osmotic Characteristics, Water Permeability, and Cryoprotectant Permeability of Human Vaginal Immune Cells. Cryobiology. 2016; doi:10.1016/j.cryobiol.2016.03.003

5. Shu Z, Hughes SM, Fang C, Hou Z, Zhao G, Fialkow M, et al. Determination of the Membrane Permeability to Water of Human Vaginal Mucosal Immune Cells at Subzero Temperatures Using Differential Scanning Calorimetry. Biopreserv Biobank. 2016; bio.2015.0079. doi:10.1089/bio.2015.0079

6. McKinnon LR, Hughes SM, De Rosa SC, Martinson JA, Plants J, Brady KE, et al. Optimizing viable leukocyte sampling from the female genital tract for clinical trials: an international multisite study. Tachedjian G, editor. PLoS One. 2014;9: e85675. doi:10.1371/journal.pone.0085675

7. Critchfield JW, Lemongello D, Walker DH, Garcia JC, Asmuth DM, Pollard RB, et al. Multifunctional human immunodeficiency virus (HIV) gag-specific CD8+ T-cell responses in rectal mucosa and peripheral blood mononuclear cells during chronic HIV type 1 infection. J Virol. 2007;81: 5460–71. doi:10.1128/JVI.02535-06

8. R Core Team. R: A Language and Environment for Statistical Computing [Internet]. Vienna, Austria; 2015. Available: http://www.r-project.org/

9. Wickham H, Francois R. dplyr: A Grammar of Data Manipulation [Internet]. 2015. Available: http://cran.r-project.org/package=dplyr

10. Bache SM, Wickham H. magrittr: A Forward-Pipe Operator for R [Internet]. 2014. Available: http://cran.r-project.org/package=magrittr

11. Wickham H. Reshaping Data with the reshape Package. J Stat Softw. 2007;21: 1–20. Available: http://www.jstatsoft.org/v21/i12/

12. Wickham H. tidyr: Easily Tidy Data with ‛spread()‛ and ‛gather()‛ Functions [Internet]. 2016. Available: https://cran.r-project.org/package=tidyr

13. Wickham H, Francois R. readr: Read Tabular Data [Internet]. 2015. Available: https://cran.r-project.org/package=readr

14. Neuwirth E. RColorBrewer: ColorBrewer Palettes [Internet]. 2014. Available: http://cran.r-project.org/package=RColorBrewer

15. Brewer CA. Color Brewer [Internet]. 2015 [cited 8 Feb 2016]. Available: http://www.colorbrewer.org

16. Pinheiro J, Bates D, DebRoy S, Sarkar D, R Core Team. nlme: Linear and Nonlinear Mixed Effects Models [Internet]. 2015. Available: http://cran.r-project.org/package=nlme

17. Hothorn T, Bretz F, Westfall P. Simultaneous Inference in General Parametric Models. Biometrical J. 2008;50: 346–363.

18. Harrell Jr FE. Hmisc: Harrell Miscellaneous [Internet]. 2015. Available: http://cran.r-project.org/package=Hmisc

19. Wickham H. ggplot2: Elegant Graphics for Data Analysis [Internet]. Springer-Verlag New York; 2009. Available: http://had.co.nz/ggplot2/book

20. Daroczi G, Tsegelskyi R. pander: An R Pandoc Writer [Internet]. 2015. Available: http://cran.r-project.org/package=pander

21. Xie Y. knitr: A General-Purpose Package for Dynamic Report Generation in R [Internet]. 2015. Available: http://yihui.name/knitr/

22. Xie Y. Dynamic Documents with R and knitr [Internet]. 2nd ed. Boca Raton, Florida: Chapman and Hall/CRC; 2015. Available: http://yihui.name/knitr/

23. Lee Y-A, Kim Y-H, Kim B-J, Kim B-G, Kim K-J, Auh J-H, et al. Cryopreservation in trehalose preserves functional capacity of murine spermatogonial stem cells. PLoS One. 2013;8: e54889. doi:10.1371/journal.pone.0054889

24. Sasnoor LM, Kale VP, Limaye LS. Supplementation of conventional freezing medium with a combination of catalase and trehalose results in better protection of surface molecules and functionality of hematopoietic cells. J Hematother Stem Cell Res. 2003;12: 553–64. doi:10.1089/152581603322448268

25. Fuller BJ. Cryoprotectants: the essential antifreezes to protect life in the frozen state. Cryo Letters. 25: 375–88. Available: http://www.ncbi.nlm.nih.gov/pubmed/15660165

26. Imaizumi K, Nishishita N, Muramatsu M, Yamamoto T, Takenaka C, Kawamata S, et al. A simple and highly effective method for slow-freezing human pluripotent stem cells using dimethyl sulfoxide, hydroxyethyl starch and ethylene glycol. PLoS One. 2014;9: e88696. doi:10.1371/journal.pone.0088696

27. Germann A, Schulz JC, Kemp-Kamke B, Zimmermann H, von Briesen H. Standardized Serum-Free Cryomedia Maintain Peripheral Blood Mononuclear Cell Viability, Recovery, and Antigen-Specific T-Cell Response Compared to Fetal Calf Serum-Based Medium. Biopreserv Biobank. 2011;9: 229–236. doi:10.1089/bio.2010.0033

28. Cohen CR, Moscicki A-BA, Scott MEM, Ma Y, Shiboski S, Bukusi E, et al. Increased levels of immune activation in the genital tract of healthy young women from sub-Saharan Africa. AIDS (London,. 2010;24: 2069–74. doi:10.1097/QAD.0b013e32833c323b

29. Muyanja E, Ssemaganda A, Ngauv P, Cubas R, Perrin H, Srinivasan D, et al. Immune activation alters cellular and humoral responses to yellow fever 17D vaccine. J Clin Invest. 2014;124: 3147–3158. doi:10.1172/JCI75429

30. Filbert H, Attig S, Bidmon N, Renard BY, Janetzki S, Sahin U, et al. Serum-free freezing media support high cell quality and excellent ELISPOT assay performance across a wide variety of different assay protocols. Cancer Immunol Immunother. 2013;62: 615–627. doi:10.1007/s00262-012-1359-5

31. Schulz JC, Germann A, Kemp-Kamke B, Mazzotta A, von Briesen H, Zimmermann H. Towards a xeno-free and fully chemically defined cryopreservation medium for maintaining viability, recovery, and antigen-specific functionality of PBMC during long-term storage. J Immunol Methods. Elsevier B.V.; 2012;382: 24–31. doi:10.1016/j.jim.2012.05.001

32. Ducar C, Smith D, Pinzon C, Stirewalt M, Cooper C, McElrath MJ, et al. Benefits of a comprehensive quality program for cryopreserved PBMC covering 28 clinical trials sites utilizing an integrated, analytical web-based portal. J Immunol Methods. 2014;409: 9–20. doi:10.1016/j.jim.2014.03.024

33. Stroncek DF, Xing L, Chau Q, Zia N, McKelvy A, Pracht L, et al. Stability of cryopreserved white blood cells (WBCs) prepared for donor WBC infusions. Transfusion. 2011;51: 2647–55. doi:10.1111/j.1537-2995.2011.03210.x

34. Veeraputhiran M, Theus JW, Pesek G, Barlogie B, Cottler-Fox M. Viability and engraftment of hematopoietic progenitor cells after long-term cryopreservation: effect of diagnosis and percentage dimethyl sulfoxide concentration. Cytotherapy. 2010;12: 764–766. doi:10.3109/14653241003745896

